# Disentangling presentation and processing times in the brain

**DOI:** 10.1101/566042

**Authors:** Laurent Caplette, Robin A. A. Ince, Karim Jerbi, Frédéric Gosselin

## Abstract

Visual object recognition seems to occur almost instantaneously. However, not only does it require hundreds of milliseconds of processing, but our eyes also typically fixate the object for hundreds of milliseconds. Consequently, information reaching our eyes at different moments is processed in the brain together. Moreover, information received at different moments during fixation is likely to be processed differently, notably because different features might be selectively attended at different moments. Here, we introduce a novel reverse correlation paradigm that allows us to uncover with millisecond precision the processing time course of specific information received on the retina at specific moments. Using faces as stimuli, we observed that processing at several electrodes and latencies was different depending on the moment at which information was received. Some of these variations were caused by a disruption occurring 160-200 ms after the face onset, suggesting a role of the N170 ERP component in gating information processing; others hinted at temporal compression and integration mechanisms. Importantly, the observed differences were not explained by simple adaptation or repetition priming, they were modulated by the task, and they were correlated with differences in behavior. These results suggest that top-down routines of information sampling are applied to the continuous visual input, even within a single eye fixation.

## 1 Introduction

Visual object recognition is a process that seems to occur almost instantaneously. However, this is just an impression: not only does our brain process the object for hundreds of milliseconds, but we will typically fixate it for hundreds of milliseconds too. Of course, light reflected on the object continually hits our retina throughout this fixation. The light reaching our eyes at each specific moment will then be processed in the brain. Since processing takes some time, light reaching our eyes at different moments during the fixation will typically be processed in the brain at the same moment (but possibly at different processing levels; Figure 1). The brain activity evoked by the perception of an object is a combination of the brain responses to information received on the retina at different moments.

**Figure 1.**
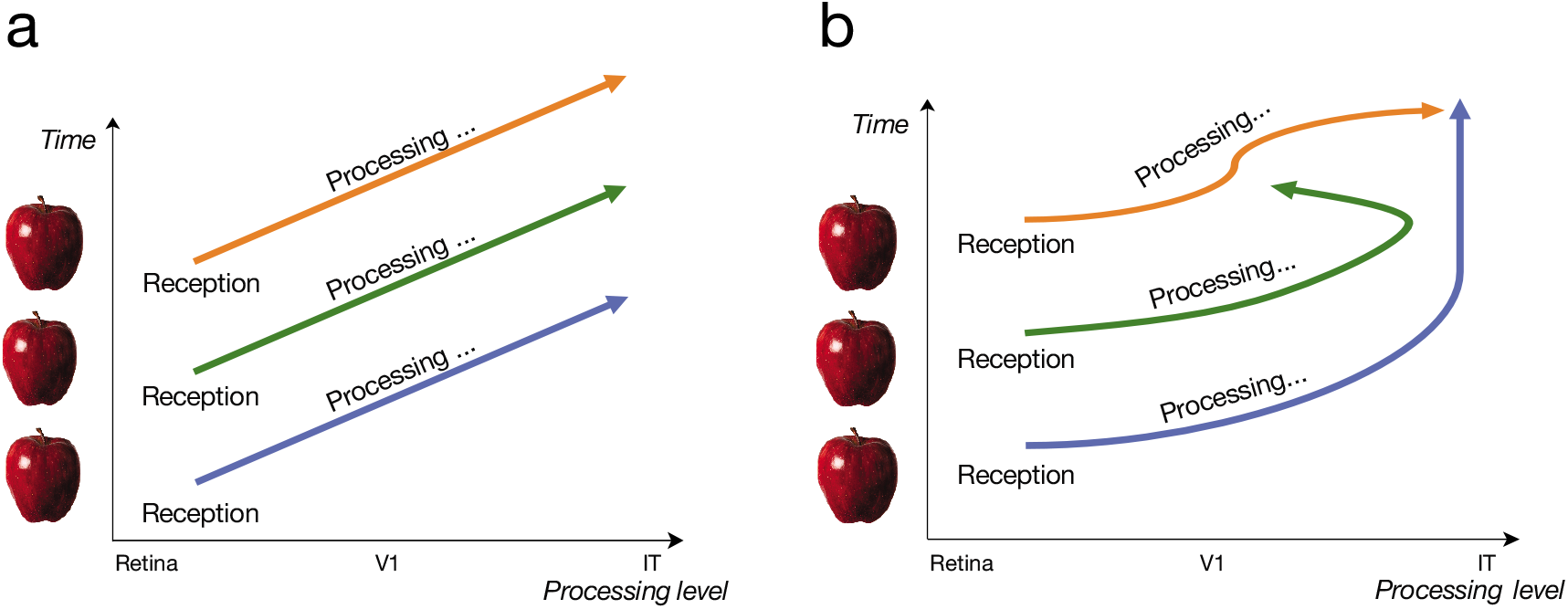
At any given point in time (any horizontal imaginary line in the above graphs), information received at different moments during fixation is simultaneously processed in the brain (possibly at different processing levels). **A)** Processing is identical for information received at different moments. **B)** Processing is different for information received at different moments.

We can expect visual information received at different moments to be processed differently (Figure 1b). This is partly because of the limited processing capacity of higher visual areas^1–2^, which prevents too much information from being processed simultaneously. One strategy that can be applied by the visual system to overcome this limitation is to use visual information received in different time windows to process different features (e.g., different regions of space, colors or spatial frequencies). This is often referred to as topdown attention being guided from one feature to another^3–4^, as a visual routine^5^, or simply as a sampling of different features across time.

The use of the information received at specific moments to process specific features may arise because this is a more efficient strategy for some tasks than using information received at any moment to process any feature^5^. Moreover, specific strategies may be more efficient than others. For example, it may be computationally more efficient to process coarse information before finer noisier features, when recognizing objects or scenes^6–7^, and so, high visual areas might process coarse information received early and fine information received late but not fine information received early. It follows that relatively stable strategies may occur in individuals, or even across individuals. Other biases may also result in stable strategies: for example, a tendency to process the most informative features in the information received first (which is probably an evolutionarily sensible strategy), or an attempt to compensate anatomical limitations (e.g., process color from the information received earlier because color is processed more slowly^8–9^). These strategies are likely to depend on the expected input and on the task.

How information received at different moments within a fixation is processed for object recognition is rarely investigated, possibly in part because the distinction between stimulus presentation time and processing time is not often discussed or appreciated (but see 10). Still, a few behavioral studies have examined this question, either by randomly revealing image features across time^9,11–14^ or by adding noise that is randomly varying across time^15–16^, and by correlating the samples with the subject’s response. These methods and similar ones (e.g., randomly varying inter-stimulus intervals with high resolution) have been employed several times in the related literature on attention and detection mechanisms^17–22^. Using such methods in object recognition paradigms has led to multiple demonstrations of how observers use the information received at different moments to categorize an object. Interestingly, these strategies often seem stable across individuals. For example, as it was hypothesized, correct responses correlate with high spatial frequency, or fine, information received late, and with low spatial frequency, or coarse, information received early and late^12–13,23^ (see also 24-25). These strategies also seem to be contingent on the task at hand^26^.

While studies have been conducted on the effects of stimulus onset asynchrony^27^, duration^28–29^, and ordering^30^ on brain activity, the processing by the brain of information received at specific *moments* during a fixation has, to our knowledge, never been investigated. This a fundamentally different endeavor: decomposing the processing time course of an object according to the moment at which information is received should inform us about the neural mechanisms underlying the differential sampling and integration of information across time. It should allow us to disentangle the sampling and the processing of visual information, which are both unraveling through time.

In this study, we aimed to perform such a decomposition. To do so, we randomly sampled the features of a face across time while subjects were performing a gender or expression recognition task^9,14^ (Figure 2; Movies S1-S4) and while their EEG activity was recorded. Faces were chosen as stimuli because they are important social stimuli that human brains are wired by evolutionary pressures to process efficiently; moreover, faces are particularly well suited to a spatial sampling of information as they all are composed of the same spatial features with essentially the same spatial configuration. To ensure that subjects could initiate a potential top-down sampling strategy on time, face stimuli occurred at predictable moments. We then reverse correlated brain activity at all time points to information presented in different time windows. We had three main hypotheses: 1) the processing time course of information received at different moments will be different; 2) this modulation of processing by the time at which information is received will itself be modulated by the task; and 3) variations in the processing of information received at different moments will correlate to variations in the behavioral use of this information for the task.

**Figure 2.**
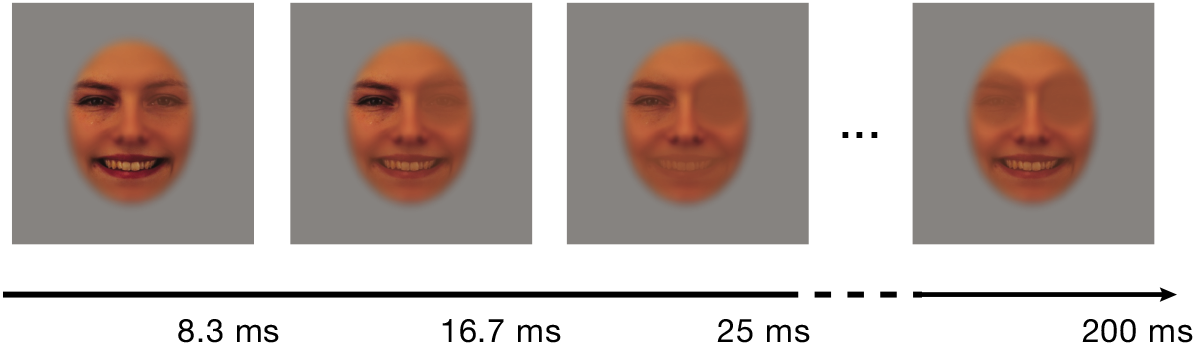
Example of a video stimulus used in a random trial. The three face features were smoothly revealed in random frames (1 frame each 8.3 ms) across 200 ms. See movies S1-S4.

## 2 Materials and Methods

### 2.1 Participants

Twenty-four neurotypical adults (mean age = 23.0 years; SD = 2.9) were recruited on the campus of the University of Montreal. Participants did not suffer from any psychiatric or psychological disorder and had no known history of head concussions. The experimental protocol was approved by the ethics board of the Faculty of Arts and Sciences of the University of Montreal and the study was carried in accordance with the approved guidelines. Written informed consent was obtained from all the participants after the procedure had been fully explained, and a monetary compensation was provided upon completion of each experimental session.

### 2.2 Materials

The experimental program ran on a Ciara Discovery computer with Windows 7 in the Matlab environment, using custom scripts and functions from the Psychophysics Toolbox^42–44^. Stimuli were shown on an Asus VG278H monitor, calibrated to allow a linear manipulation of luminance, with a resolution of 1920 × 1080 pixels and a 120 Hz refresh rate. Luminance values ranged from 2.47 cd/m^2^ to 269 cd/m^2^. A chin rest was used to maintain a viewing distance of 76 cm. EEG activity was recorded using an ANT Neuro Waveguard 64-electrode cap with Ag/AgCl electrodes, using a sampling rate of 1024 Hz and a resolution of 12 bits. Linked mastoids served as initial common reference. Vertical electro-oculogram (vEOG) was bipolarly registered above and below the dominant eye and horizontal electro-oculogram (hEOG) at the outer canthi of both eyes. Electrode impedance was kept below 10 kW during recording.

### 2.3 Stimuli and sampling

Two hundred and sixty-four color images of faces were selected from the image database *Karolinska Directed Emotional Faces* (KDEF)^45^; only faces facing the camera were chosen. These were composed of 66 different identities (33 women and 33 men) each performing a happy and a neutral expression; two different pictures of each facial expression were used. Faces were aligned on twenty hand-annotated landmarks averaged to six mean coordinates for left and right eyes, left and right eyebrows, nose and mouth, using a Procrustes transformation.

We then created an uninformative face background by taking the mean of all aligned faces and applying a lightly smoothed elliptical mask (horizontal radius = 6 degrees of visual angle) to conceal the background, hair and shoulders. The areas including and surrounding the eyes and eyebrows were then covered by two lightly smoothed approximately circular masks; the area including and surrounding the mouth was covered by a lightly smoothed elliptical mask. The color of these masks was the mean color of the unmasked parts of the average face. The three feature masks were of equal area (within a <1% margin; since feature masks were smoothed, area covered was computed by summing the mask pixel values).

For use in the sampled-face trials, the mean luminance and the contrast of all aligned faces (within the feature areas determined by the feature masks previously discussed) were equalized, separately for each color channel, using the SHINE toolbox^46^. The same procedure was applied but for the whole face (inside the elliptical mask), for use in the whole-face trials.

On each sampled-face trial, the face features of a randomly selected exemplar face were gradually revealed at random moments across a total duration of 200 ms; that is, masked feature areas of the uninformative face background were replaced by the features of an exemplar face (Figure 2; Movies S1-S4). A duration of 200 ms was chosen so that no saccade would occur during stimulus presentation on most trials. Specifically, on each trial, a random 3 × 72 sparse matrix composed of zeros and a few ones (the probability of each element being one was constant and was 0.025) was created; each row of 72 elements was then convolved with a 1-D gaussian kernel, or “bubble”^14,32^, with a 1.8 frame (15 ms) standard deviation. Superfluous padding was removed so that the final smoothed matrix was 3 × 24 in size and thresholding was applied so that no value exceeded 1. We called this matrix *sampling matrix* and the value of each element determined the visibility of a given face feature through the feature background in a given video frame for this trial; more precisely, ***P**_ijk_* = ***f**_ij_* · *s_ijk_* + *b* · (1 – *s_ijk_*), where ***p**_ijk_* are the pixel values to be displayed for face feature *i* on frame *j* in trial *k*, ***f**_ik_* are the original pixel values of face feature *i* of the exemplar face selected for trial *k*, *s_ijk_* is the sampling matrix value for face feature *i* on frame *j* in trial *k*, and *b* is the feature background color.

### 2.4 Experimental design

Each participant came to the laboratory twice and filled in a personal information questionnaire (education, age, sex, hours of sleep, alertness, concussion history, mental illness history, etc.) on the first session. Participants completed a total of 1000 sampled-face trials in each session; nine participants also completed in each session 100 additional whole-face trials in which a non-sampled exemplar face was shown for the same amount of time. Sampled-face and whole-face trials were randomly intermixed throughout the experiment. Each experimental session was divided in four equal-size blocks (of 250 or 275 trials) and blocks were interleaved with breaks of approximately 5 minutes. In addition, after every 5 trials, the screen automatically showed text indicating that the participants could take a few seconds to blink and rest their eyes before pressing a key to continue the experiment (participants were instructed not to blink during the trials themselves).

On each trial, a central fixation cross was shown to the participants for 1500 ms, after which the video stimulus appeared during 200 ms, superposed to the fixation cross, again followed by the fixation cross until the participant responded (the next trial then followed after an additional constant 1500 ms); a mid-gray background was always present. A fixed inter-trial interval was used so that participants could predict the onset of the trials. Half of the participants had to categorize the sex of the faces while the other half had to categorize their expression (happy or neutral). Participants had to respond as accurately and rapidly as possible with two keys on the keyboard (half of the participants had to use the opposite key combination from the other half, to counterbalance any motor effect).

### 2.5 Behavioral data analysis

One session from one participant was removed from all analyses because its mean accuracy was 50%; a session from a different participant was removed because of prominent EEG artifacts on a large subset of trials. Finally, one 275-trial block from still another participant was lost due to a technical error.

Accuracies and response times were z-scored within each 250- or 275-trial block. Trials with a z-scored response time below −3 or above 3, or with an absolute response time below 100 ms or above 2000 ms, were excluded from further analyses. Sampling matrices weighted by z-scored accuracies were then averaged together for each session. (Such a weighted sum is equivalent to a linear regression here since sampling was random.) Resulting *classification images* were averaged together within each subject and then within each task. Analyses were repeated with randomly permuted accuracies 10,000 times and a statistical threshold (*p* < .05, one-tailed, pixel level, corrected for familywise error rate (FWER)) was determined using the maximum statistic method^47^. Since we were only interested in which information was used to do the task, we only assessed positive correlations and performed a one-tailed test.

### 2.6 EEG data preprocessing

All preprocessing was performed with the help of functions from the Fieldtrip toolbox^48^. EEG raw data from each session was segmented in trials, filtered between 1 and 30 Hz with two successive 4^th^ order Butterworth IIR filters, baseline corrected using the average activity between 500 ms and 250 ms before stimulus presentation, and downsampled to a 250 Hz sampling rate. Mastoid electrodes were removed due to poor signal-to-noise ratio on most subjects and data was re-referenced to an average reference. Anomalous trials, trials in which eye movements were occurring during the stimulus and anomalous electrodes were identified and removed following careful visual inspection of the data (mean number of trials = 4.5 (0.5%), SD = 9.22 (0.9%)); bad channels were interpolated using a spherical spline (mean number of channels = 1.02, SD = 0.81). An ICA using Hyvärinen’s fixed-point algorithm^49^ was then performed to identify blink and eye movement artifacts. Bad components were identified and removed following careful visual inspection (mean number of components = 1.38, SD = 0.65). Finally, we computed single-trial current scalp density (CSD) waveforms using the spherical spline method (lambda = 1e-5, spline order = 4, degree of Legendre polynomials = 14)^50–51^; all further analyses were conducted on this CSD data.

### 2.7 EEG data analysis

#### 2.7.1 Falsely correct trials

In every experiment in which performance is not at ceiling level, part of the trials initially labeled as correct are correct only by chance: e.g., if 20% of responses are incorrect, this means that another 20% was in fact correct only by chance (since there is a 50% chance of being correct or incorrect when guessing). Here, we can verify which trials are comprised in this percentage of “falsely” correct trials by verifying which are the trials whose sampling matrices correlate the least to the behavioral classification image. Using this novel analysis method, we kept only true correct trials which were not correct merely by chance for further analyses.

#### 2.7.2 Regression analyses

Trials with a z-scored response time below −3 or above 3, or with an absolute response time below 100 ms or above 2000 ms, were excluded from the regression analyses. For each session, electrode and time point, regularized (ridge) multiple linear regressions were performed between the standardized feature × presentation time sampling planes and the standardized EEG amplitudes (Figure S1a). Resulting regression coefficients were convolved with a Gaussian kernel (standard deviation of 3 time points, or 12 ms) in the EEG time dimension. Maps of regression coefficients were averaged within each subject and then across subjects within each task. Analyses were repeated with randomly permuted trials 1,000 times and statistical thresholds (*p* < .05, two-tailed, FWER-corrected) at both the pixel and cluster (2D clusters across EEG time and presentation time; using the summed cluster values; arbitrary primary threshold of *p* < .01, two-tailed, uncorrected) levels were determined using the maximum statistic method^47^. Analyses were restricted to time points between 30 ms and 600 ms from face onset. Results are displayed for representative PO7 (left occipito-temporal; LOT) and PO8 (right occipito-temporal; ROT) sensors but multiple comparison corrections were applied across all electrodes. Results were similar for most occipito-temporal sensors; data from all electrodes is available in an online repository (https://osf.io/3r782/).

#### 2.7.3 Task × stimulus moment ANOVA

To investigate whether processing was significantly modulated by the presentation moment and the task, a task × presentation moment ANOVA was performed. Maps of regression coefficients for each subject, face feature and electrode were first linearly interpolated to a resolution of 0.1 ms, realigned to the feature onset instead of the face onset (e.g., the EEG activity for the first presentation moment stayed the same, while activity for the second one was shifted left by 8.3 ms, activity for the third one by 16.7 ms, and so on), and resampled to the original resolution of 4 ms. Task × presentation moment ANOVAs were then performed on individual subjects’ regression coefficients for each face feature, electrode, and latency from the feature onset (Figure S1b). Resulting F values were interpolated in topography space using biharmonic spline interpolation^52^. Analyses were repeated on the 1,000 null maps obtained by randomly permuting trials and statistical thresholds (*p* < .05, one-tailed, FWER-corrected) at both the pixel and cluster (3D clusters across EEG time and topography space; using the summed cluster values; arbitrary primary threshold of *p* < .01, one-tailed, uncorrected) levels were determined using the maximum statistic method^47^. A one-tailed test was performed given that F statistics are non-negative. Analyses were restricted to time points between 50 ms and 400 ms from feature onset.

### 2.8 Mutual information between brain and behavior regression coefficients

For each subject, electrode and latency from feature onset, Gaussian copula mutual information^53–54^ was computed between the results of the behavior-stimulus weighted sum and the absolute values of the results of the EEG-stimulus regression, across stimulus moments (stimulus presentation time frames). Analyses were repeated with regression coefficients from the 1,000 null maps obtained by randomly permuting trials and statistical thresholds (*p* < .05, one-tailed, FWER-corrected) at both the pixel and cluster (3D clusters across EEG time and topography space; using the summed cluster values; arbitrary primary threshold of *p* < .01, one-tailed, uncorrected) levels were determined using the maximum statistic method^47^. A one-tailed test was performed given that mutual information is nonnegative. Analyses were restricted to time points between 50 ms and 400 ms from feature onset.

## 3 Results

### 3.1 Time course of information use

Mean accuracy was 75.8% (s = 4.2%) in the gender task and 82.9% (s = 6.2%) in the expression task. Mean response time was 711 ms (s = 87 ms) in the gender task and 662 ms (s = 100 ms) in the expression task.

To identify which face features in which time frames led to accurate responses, we performed for each session a sum of sampling matrices (indicating the visibility of each face feature at each time frame in the stimulus on each trial) weighted by accuracies. Mean results for each task are displayed in Figure 3. As we can see, both eyes were used at all except the earliest moments, while the mouth was used throughout the presentation to identify the expression of the face. These results replicate previous studies using a spatial sampling of the whole face^9,31–33^.

**Figure 3.**
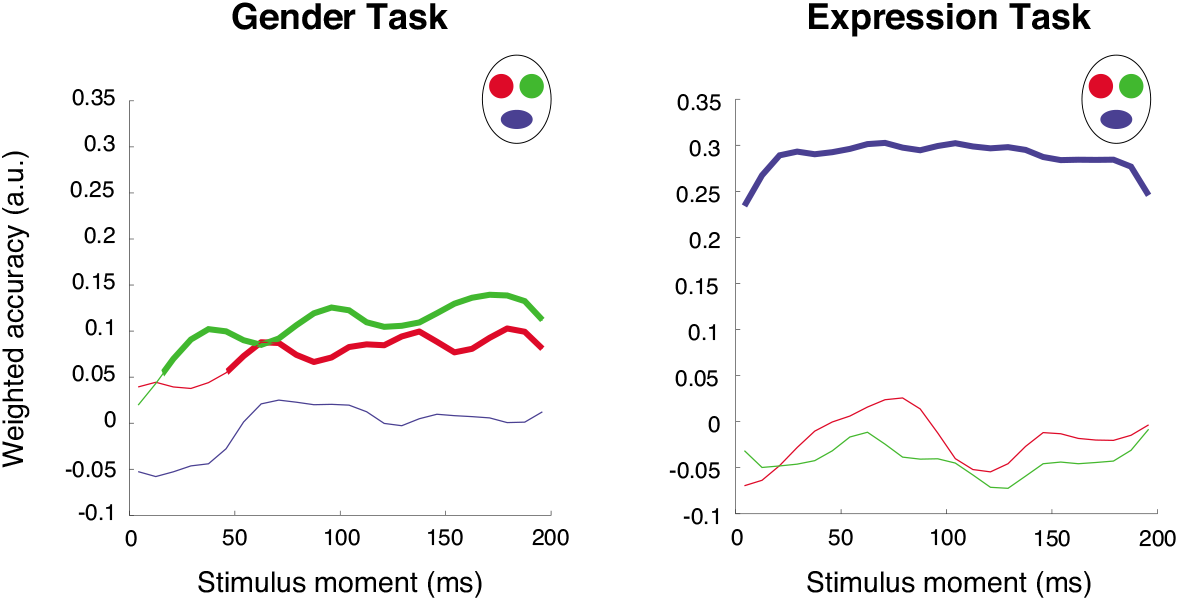
Behavioral results indicating, for each task, how each feature presented on each frame correlates with correct responses. Bold segments of line indicate frames that are significant (*p* < .05, one-tailed, FWER- corrected).

Note that these time points refer to the moment of presentation of the feature within the stimulus, and so, equivalently, to the moment at which information is received on the retina. To avoid any confusion with processing time (as assessed with EEG), we refer to this time dimension as stimulus time; to avoid any confusion with stimulus duration, we will usually refer to stimulus “moments”.

### 3.2 Visual Evoked Potentials

To verify if our sampling method elicited, on average, similar ERPs to whole unaltered faces, we computed the average of all trials with sampled and whole faces, for those subjects who performed the task on both kinds of trials. We display the ERPs of representative left and right occipito-temporal sensors (LOT and ROT), and the overall topographies (Figure 4). As we can see, ERPs and their associated topographies are very similar between the conditions. We computed the difference between the ERPs and assessed its significance using a paired permutation test (500 permutations): there was no significant difference between the conditions at any time point on either sensor (*p* > .05, two-tailed, FWER-corrected with the maximum statistic method). This suggests that our sampling method did not greatly alter the average brain response to faces.

**Figure 4.**
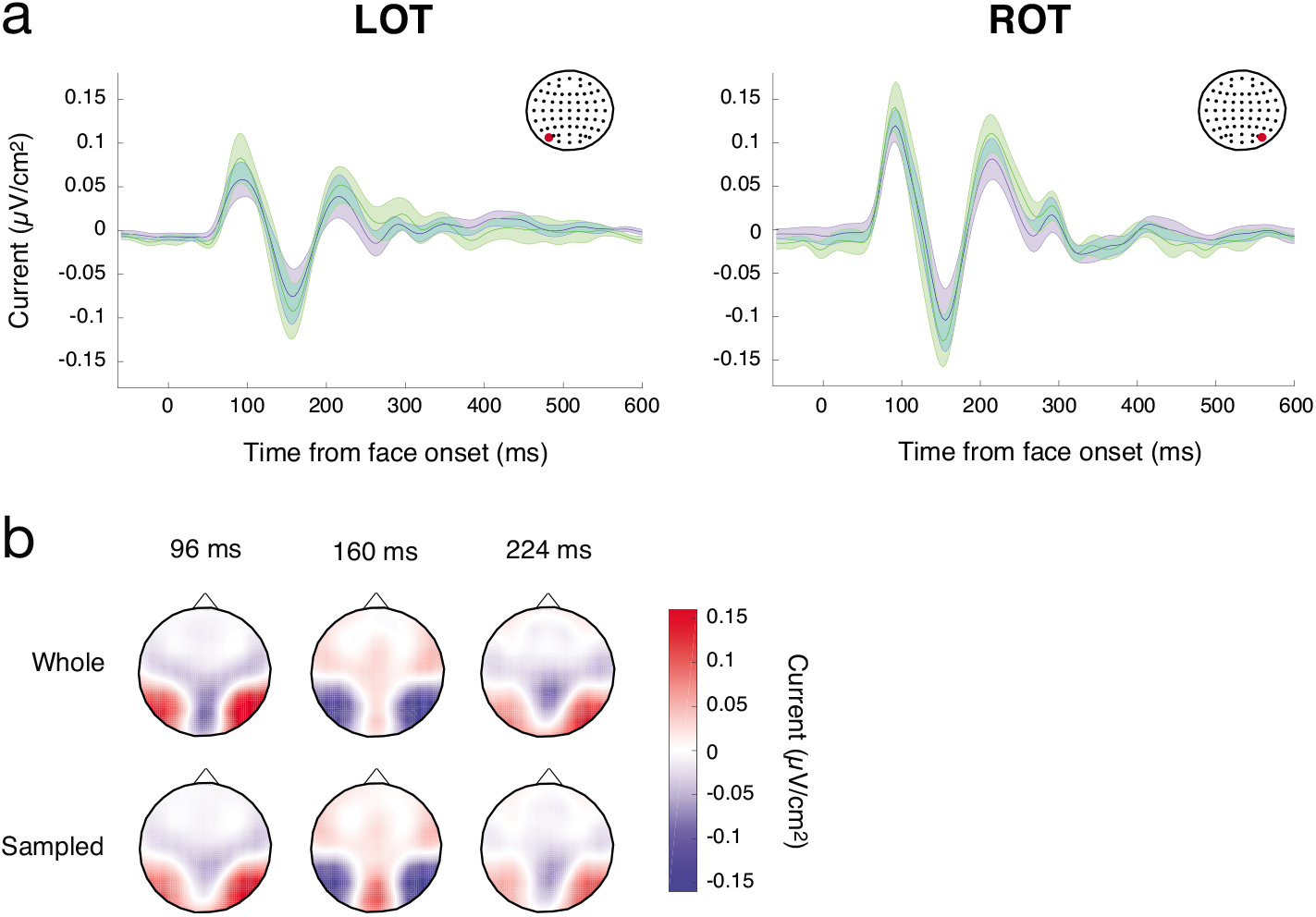
Mean ERPs for whole (green) and sampled (blue) faces on LOT and ROT. **A)** Shaded areas represent standard errors above and below the mean. **B)** Topographies for whole and sampled faces at selected latencies.

### 3.3 Uncovering the processing of information received at different moments

For each session, ridge regressions were performed between sampling matrices of correct trials and EEG amplitude on each time point and electrode (see Methods; Figure S1a). Although analyses were conducted on all electrodes (and appropriate corrections for multiple comparisons were applied), we will mostly focus on results from occipitotemporal sensors (see also Figure S4 for summary scalp maps computed using global power). Mean maps of regression coefficients are displayed for representative left and right occipito-temporal sensors (LOT and ROT) on Figures 5 (gender task) and 6 (expression task). These maps show a complete portrait of what is happening during visual recognition: how information impinging the retina at different moments throughout fixation is simultaneously processed through time in the brain.

**Figure 5.**
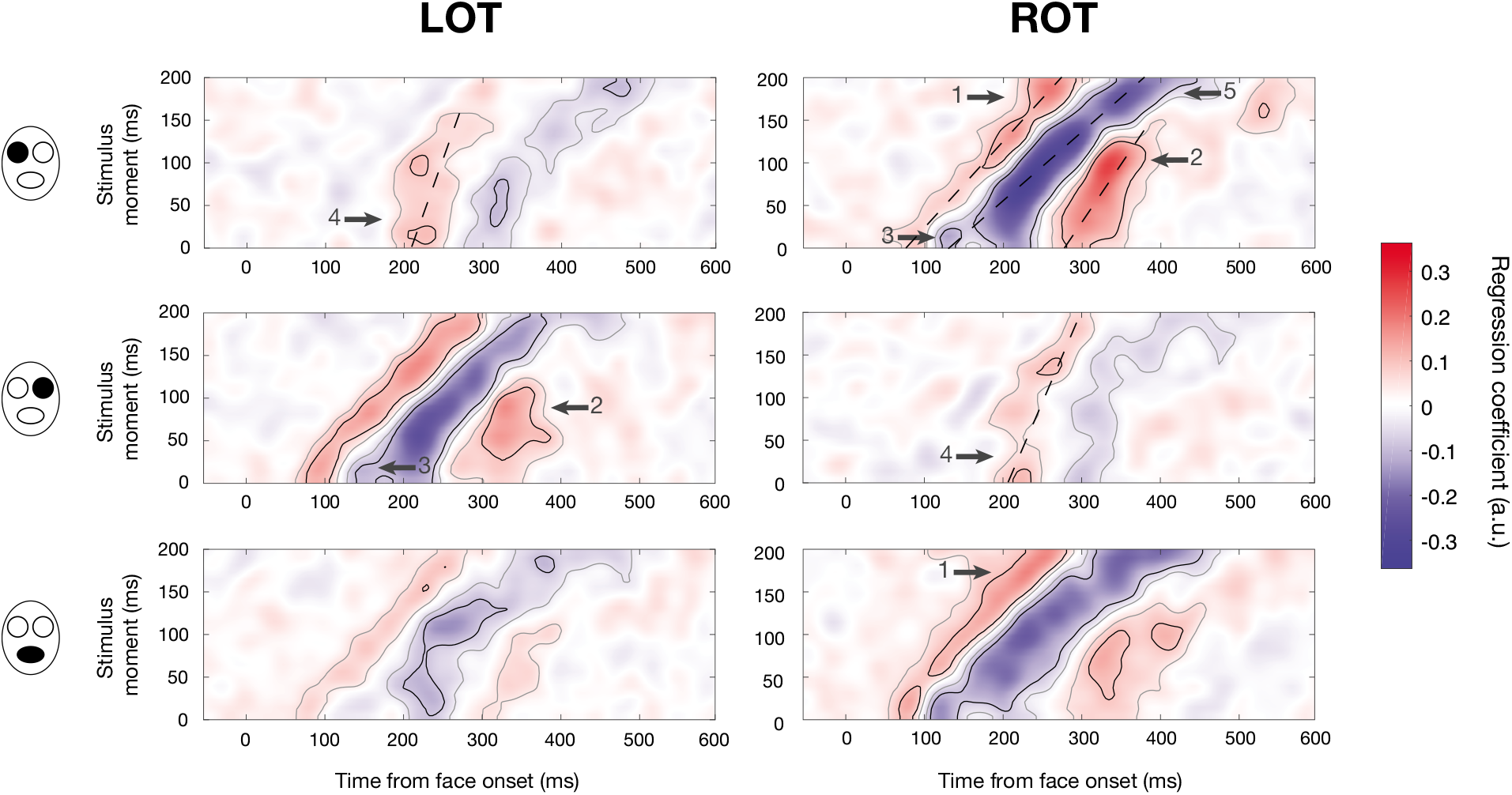
Mean maps of regression coefficients for each face feature (rows) on LOT and ROT sensors (columns) for the gender task. Within each map, each row refers to the EEG activity (across time) related to the presentation of the face feature on a given frame within the stimulus, i.e. the processing of information received on the retina at a specific moment. Gray outlines indicate significance at the cluster level and black outlines indicate significance at the pixel level (*p* < .05, two-tailed, FWER-corrected). Dashed lines illustrate components with slopes different from one. Arrows point toward some results of interest: (1) an increase in early activity for information received later; (2) late activity is maximal for information received mid-fixation; (3) additional negative peak for information received at the fixation onset; (4) large latency shift for activity related to information received early on ipsilateral electrodes; and (5) increased latency of negative activity for information received at the end of fixation.

We can immediately see on most maps (especially the ones for the mouth and the contralateral eyes) a clear diagonal trend: as it could be expected, information received on the retina *x* ms later is on average processed *x* ms later in the brain. This processing takes the form, in most cases, of a positive activation followed by a negative one and another positive one (analogous to the classic P1, N170 and P3 components). However, there also seem to be important differences in amplitude across stimulus moments. In the next section, we look at these differences in more details.

### 3.4 Investigating differences in processing across stimulus moments

To assess whether differences in processing across stimulus moments are statistically significant, we conducted a task × stimulus moment ANOVA on regression coefficients for each face feature, electrode and EEG latency, after having realigned each row of the previous maps so that the zero point on the *x* axis is the feature onset rather than the face onset (see Methods; Figure S1b).Significant modulation of processing by the stimulus moment is visible during almost all the analyzed time window (~50-360 ms; Figure 7). Differences are strongest on occipito-temporal sensors, but they are also present on central and frontal sensors, especially at higher latencies (e.g., there is a significant effect of stimulus moment peaking between 300 and 350 ms on frontal Fpz sensor).

**Figure 6.**
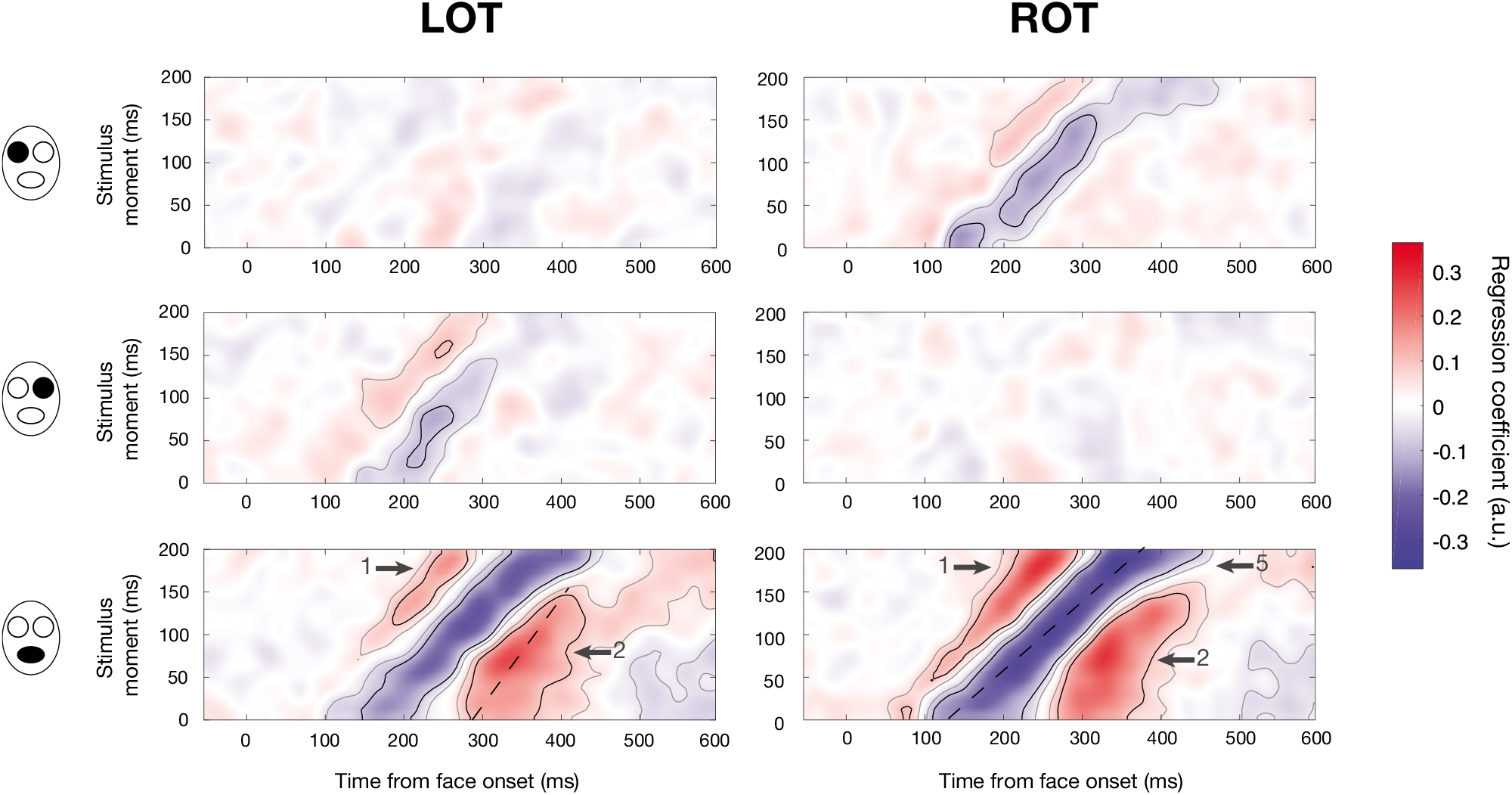
Mean maps of regression coefficients for each face feature (rows) on LOT and ROT sensors (columns) for the expression task. Within each map, each row refers to the EEG activity (across time) related to the presentation of the face feature on a given frame within the stimulus, i.e. the processing of information received on the retina at a specific moment. Gray outlines indicate significance at the cluster level and black outlines indicate significance at the pixel level (*p* < .05, two-tailed, FWER-corrected). Dashed lines illustrate components with slopes different from one. Arrows indicate results of interest : (1) an increase in early activity for information received later; (2) late activity is maximal for information received mid-fixation; (5) increased latency of negative activity for information received at the end of fixation.

**Figure 7.**
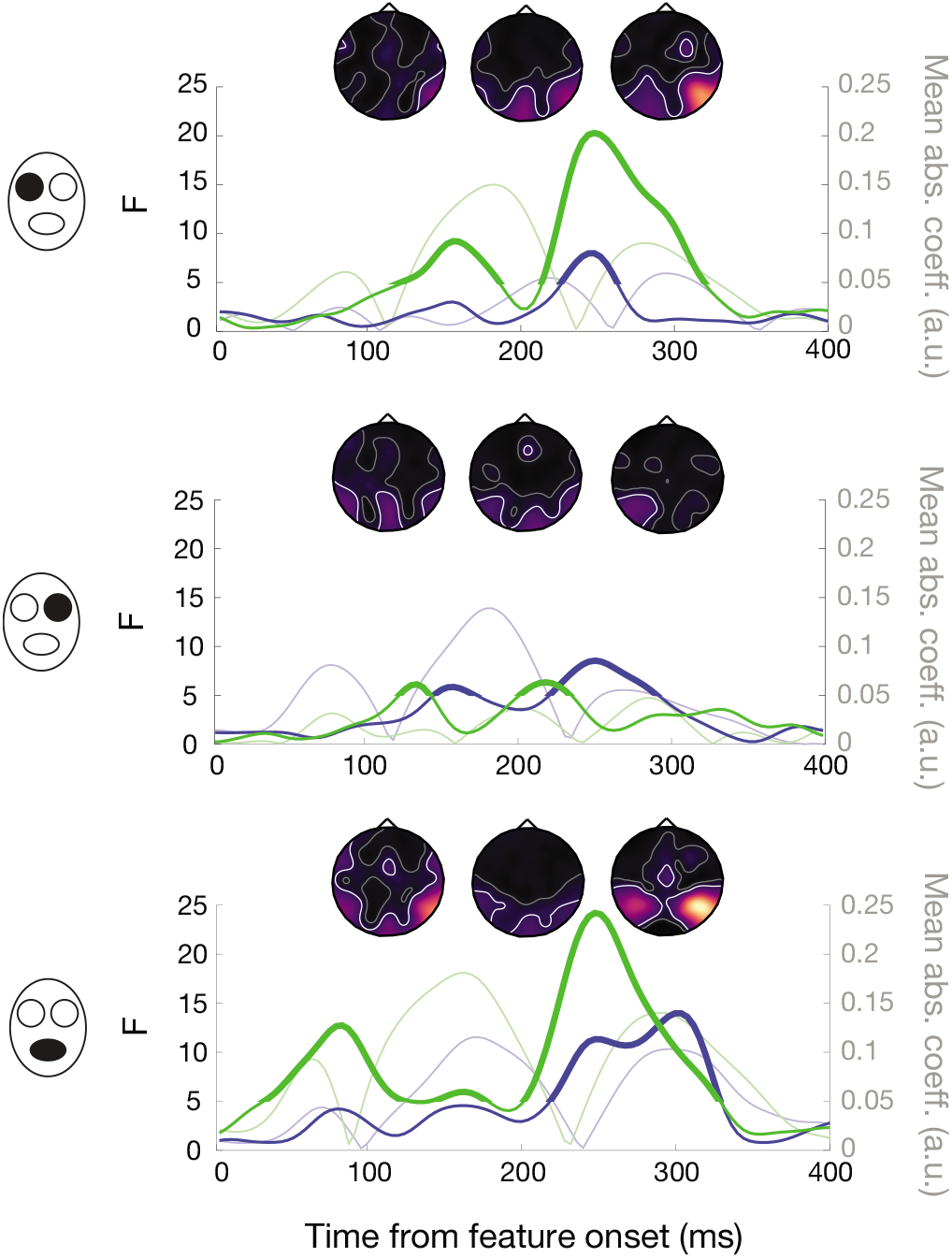
Effect of stimulus moment on EEG activity, for each face feature. F values are shown for all latencies (from the feature onset) for LOT (blue) and ROT (green) sensors; bold segments indicate time points significant at the pixel level (*p* < .05, FWER-corrected across sensors and time). These F values indicate how much activity at a given latency is influenced by the exact moment at which information is presented within the stimulus. These time courses are superposed to the mean magnitudes (across stimulus moments) of the regression coefficients (in smaller point and less saturated color). Higher F values do not necessarily coincide with higher average activity. Topographies depict the temporal progression of the effect of presentation moment across the whole scalp: latencies of 100, 150 and 250 ms are shown. Lighter colors indicate higher F values; white curves indicate areas significant at the pixel level and gray curves indicate areas significant at the cluster level (*p* < .05, one-tailed, FWER-corrected across topography and time).

On occipito-temporal sensors, variations in the amplitude of the first positive activation across stimulus moments are leading to significant differences around a latency of 80-100 ms: specifically, this activation is stronger at late stimulus moments or at all except intermediate stimulus moments (Arrow 1, Figures 5–6). The last positive activation peaking at intermediate stimulus moments is also a source of significant variations around 300 ms (Arrow 2, Figures 5–6).

#### 3.4.1 An additional negative peak for early stimulus moments

Interestingly, significant differences in amplitude around 150 ms for the contralateral eyes in the gender task are partly driven by the presence of an apparent additional peak, for the early stimulus moments (Arrow 3, Figure 5). We verified whether these two peaks represented two distinct components with different topographies. To do so, we used the maps of regression coefficients for individual sessions and looked at the topographies (one value for each electrode) associated with both peaks (at the same stimulus moment); we analyzed the 12 subjects performing the gender task. We thus had four topographies per subject and per eye: one for each peak in each session. For each subject, we computed a cosine similarity metric (1 – the absolute value of the cosine angle) between the topographies associated to the same peak on different days and averaged them: this is the within-peak similarity. Next, we computed the same metric for topographies associated to different peaks on different days and averaged them: this is the between-peaks similarity. We finally performed t-tests between these similarity metrics: the within-peak similarities were significantly greater for the right eye (*t*(11) = 4.76, *p_Bonf_* = .002) but not for the left eye (*t*(11) = 1.38, *p_Bonf_* > .10). When using the topographies associated to different peaks on the *same* day to compute the similarity metric, we still obtained significantly greater within-peak similarities for the right eye but not for the left eye (left eye: *t*(11) = 0.97, *p_Bonf_* > .10; right eye: *t*(11) = 3.07, *p_Bonf_* =.042). In other words, for the right eye feature at least, topographies associated with the same peak obtained on different days are more similar than topographies associated to different peaks, even when these are obtained on the same day. Consequently, each peak represents a distinct activation with its own topography and neural generators, with the first one being especially sensitive to the onset and stopping being receptive after only about 20 ms.

#### 3.4.2 Variations in latencies across stimulus moments

Other variations on occipito-temporal sensors seem to be driven by increases or decreases in the latency of a component across stimulus moments. To investigate this, we computed, for each major component, task and feature, the peak latency at each significant stimulus moment on LOT and ROT (significance at the cluster level; ignoring activations past 500 ms from the face onset). We then fitted a line across these latencies (see dashed lines on Figures 5 and 6) and tested (one-sample t-test) whether the slope of the line was significantly different from 1. Here, a slope of 1 would mean that the feature takes the same time to be processed at all stimulus moments, whereas a larger slope would mean that the feature takes increasingly longer to be processed with increasing stimulus moment, and a smaller slope that the feature takes an increasingly shorter time to be processed with increasing stimulus moment; a slope of 0 would mean that features are processed at the same moment irrespectively of when they were received on the retina. In most cases, the latency of the first positive component from the feature onset was approximately constant (i.e. same processing duration for all stimulus moments; slopes between 0.90 and 1.04, R^2^_adj_ > .96, df ≥ 11, *t* < 2.92, *p_Bonf_* > .10) except in the case of the right eye on LOT in the gender task, where it was slightly increasing (slope = 1.08, R^2^_adj_ = .99, *t*(22) = 3.24, *p_Bonf_* = .049) and in the case of the eyes on ipsilateral electrodes in the gender task where it was decreasing (slopes < .44, R^2^_adj_ > .27, df ≥ 17, *t* > 8.84, *p_Bonf_* < 1.2 × 10^-6^). The small slope for the eyes on ipsilateral electrodes illustrates the striking fact that this component always occurs about 220 ms after the face onset or later; information received the earliest is thus processed at about the same time as information received 50-75 ms later (Arrow 4, Figure 5). Regarding the middle negative component, its slope across stimulus moments was not different from 1 in most cases (slopes between 0.60 and 1.44, R^2^_adj_ > .45, df ≥ 16, *t* < 3.00, *p_Bonf_* > .08) except for the left eye on ROT in the gender task and for the mouth on ROT in the expression task (slopes > 1.69, R^2^_adj_ > .78, *t*(22) > 3.68, *p_Bonf_* < .02). In both these cases, the slope was significantly larger than 1. This is mostly a consequence of an increase in latency in the last stimulus moments (Arrow 5, Figures 5 and 6). Finally, in the case of the last positive component, the slope was significantly smaller than 1 for the eyes on the contralateral electrodes in the gender task and for the mouth on LOT in the expression task (slopes between 0.26 and 0.66, R^2^_adj_ > .66, df ≥ 13, *t* > 5.98, *p_Bonf_* < 2.0 × 10^-4^) and it was approximately constant for the mouth in the gender task and on ROT in the expression task (slopes = 0.67 and 0.79, R^2^_adj_ > .66, df ≥ 11, *t* < 3.45, *p_Bonf_* > .07).

### 3.5 Investigating top-down modulations

#### 3.5.1 Effect of the amount of information presented beforehand

The differences in processing across stimulus moments that we uncovered cannot be caused by differences in *what* has been seen before during a trial since sampling was random; however, *how much* was seen could have an influence, since the probability of already having shown information in a trial is greater in the last stimulus frame than in the first one. Thus, the observed differences could be caused in part by bottom-up effects such as adaptation or repetition priming. To investigate this possibility, we repeated the previous regressions only with trials in which just one bubble was revealed: despite a greatly reduced number of trials, results were remarkably similar (Pearson correlation of .95 between the maps of regression coefficients; Figures S2 and S3), suggesting that the previously observed effects are not caused by differences in the amount of information presented beforehand.

#### 3.5.2 Interaction between stimulus moment and task

The previous result alone does not completely exclude the possibility of bottom-up effects. To investigate whether differences in activity across stimulus moments could be explained at least in part by top-down mechanisms, we verified for each face feature, time point and location, whether there was a significant interaction between stimulus moment and task, i.e. if the moment at which information is received modulates processing differently depending on the task. There was a significant interaction at several time points and locations, again mostly on occipito-temporal electrodes but also in more anterior locations. Contrary to what we observed with the main effect of stimulus moment, there is almost no significant interaction around 100 ms, but the peak effects are similarly around 150 and 250 ms on right occipitotemporal sensors (Figure 8). Note that on some more anterior sensors such as CP1, significant interactions peaked after 300 ms.

**Figure 8.**
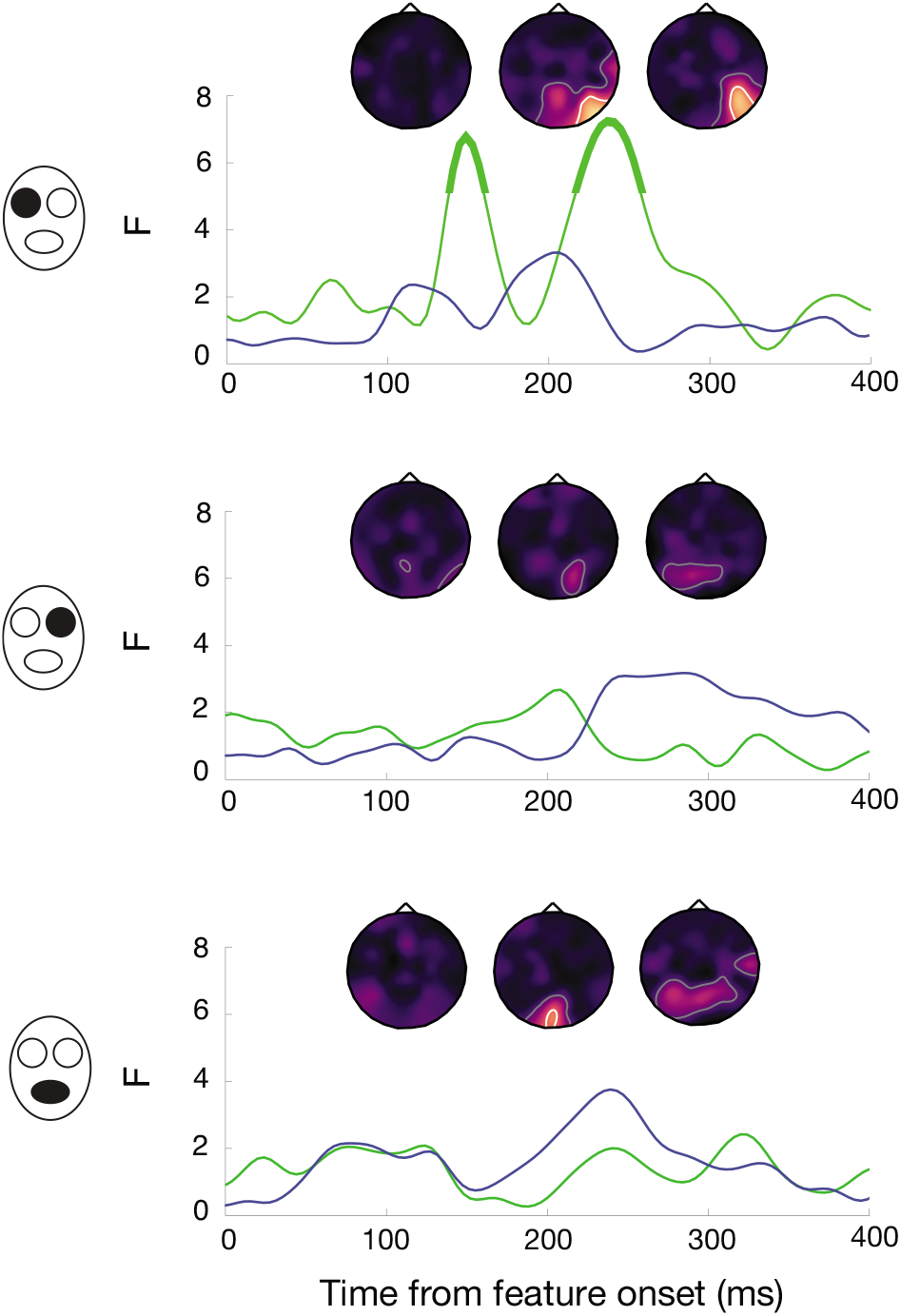
Interaction of stimulus moment and task on EEG activity, for each face feature. F values are shown for all latencies (from the feature onset) for LOT (blue) and ROT (green) sensors; bold segments indicate time points significant at the pixel level (*p* < .05, FWER-corrected across sensors and time). These F values indicate how much the activity variations across stimulus moments are influenced by the task. Topographies depict the temporal progression across the whole scalp: latencies of 100, 150 and 250 ms are shown. Lighter colors indicate higher F values; white curves indicate areas significant at the pixel level and gray curves indicate areas significant at the cluster level (*p* < .05, one-tailed, FWER-corrected across topography and time).

### 3.6 Relating sampling in the brain and in behavior

We evaluated where and when variations in brain activity across stimulus moments are related to variations in the behavioral use of information. Since differences in brain activity are likely related to the behavioral use of information in complex nonlinear ways, the mutual information (MI) metric was used. MI was computed across stimulus moments between coefficients resulting from the accuracy-weighted sums of sampling matrices (behavioral results) and the magnitudes of brain regression coefficients for each subject, face feature, latency from feature onset and electrode. Importantly, computing MI separately for each face feature allowed us to isolate the contribution of *within-feature* variations across stimulus moments. We observe significant MI mostly on occipitotemporal sensors at early and late latencies, but also in more anterior locations at later latencies (Figure 9). Regarding the eyes, significant MI is present early (<130 ms) and late (>250 ms) in both tasks, but it is present at intermediate latencies (~150-250 ms) only in the gender task. Interestingly, significant MI for the mouth is visible throughout the time course, for both tasks. While we did not uncover a significant behavioral use of the mouth in the gender task in our study, other studies have observed it, sometimes only when correlating feature visibility with response times instead of accuracy^31–33^. These results show that the origin of the variations in the use of information across stimulus moments can be traced back to variations in occipito-temporal activity at early and late latencies, and to variations in frontal activity at later latencies.

**Figure 9.**
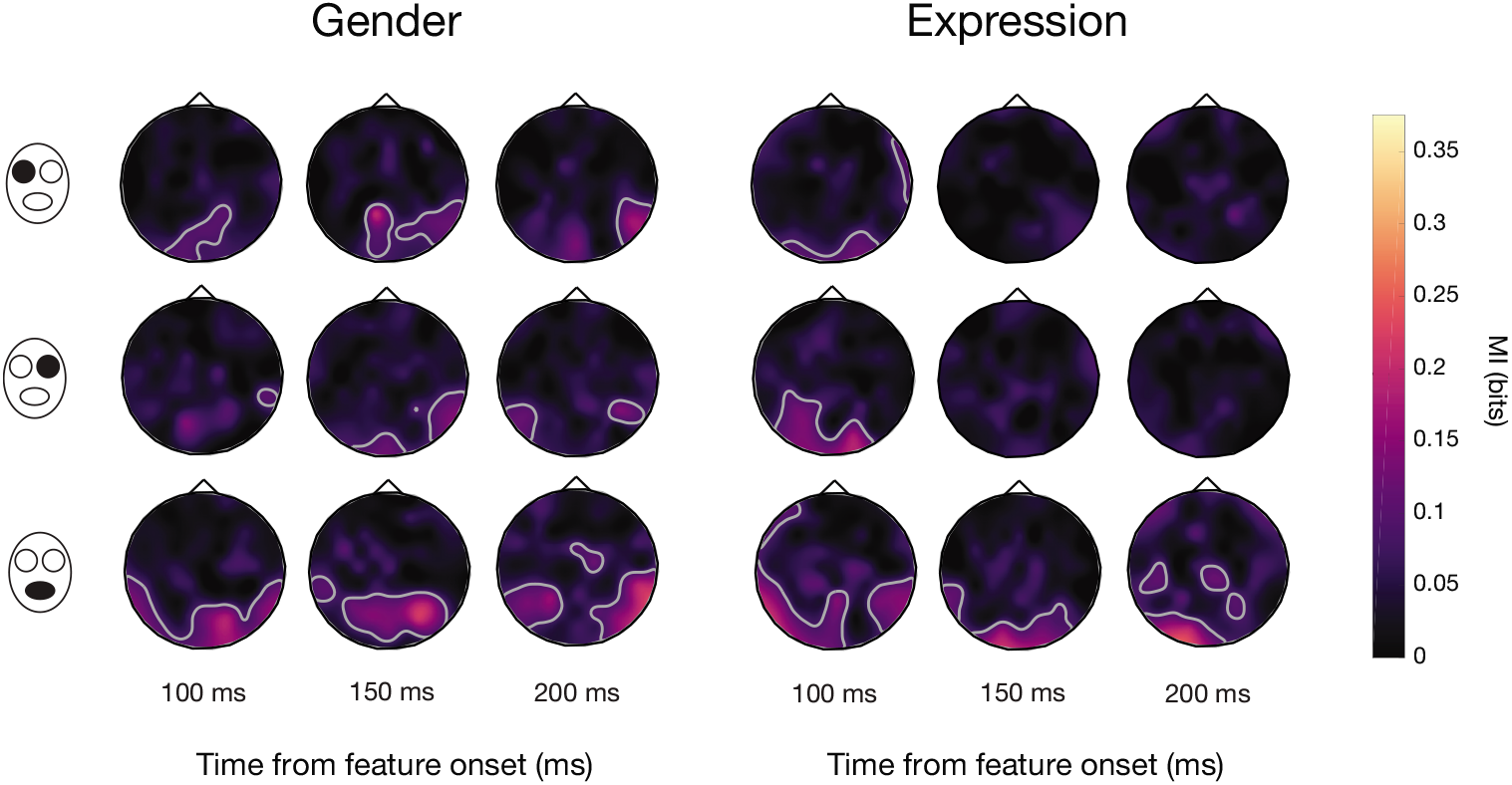
Mutual information (MI) between behavioral and brain coefficients, for selected latencies, for both tasks and all face features. High values indicate that the variations in EEG activity across stimulus moments relate to variations in behavioral accuracy across stimulus moments. Areas significant at the cluster level are outlined by gray lines (*p* < .05, one-tailed, FWER-corrected across topography and time).

## 4 Discussion

When we fixate an object, light impinges on our retinas in a continuous fashion, implying that our brain simultaneously processes information that is received at different moments, through time and cortical space. This is not typically considered in studies investigating the processing of visual objects, and so the processing uncovered in those studies corresponds to a combination of responses to information received at different moments. In our experiment, we randomly sampled the features of a face across time^14^ while brain activity was being measured to decompose this processing and uncover for the first time the brain activity related to information received at specific time points during a single eye fixation.

We first observed that information is processed differently depending on when it is received on the retina during the fixation. One of the most striking differences is seen in the ipsilateral representation of the eyes on occipito-temporal sensors in the gender task. The lateralized anatomy of the visual system tells us that each eye should be processed by the contralateral hemisphere first^34–35^: the ipsilateral representation is likely to have been transferred from an early contralateral representation^36^. Here, the contralateral representation appears to peak at a relatively constant offset of ~175 ms after information is received on the retina, independently of *when* it is received during the stimulus presentation (see the diagonal linear trend of the negative activations in Figure 5). However, the ipsilateral representation appears to be gated: all information received in the first 50 ms of fixation is represented at the same time, around 220 ms from face onset, while information received after 50 ms is represented with a fixed offset of ~120 ms, representation moment increasing linearly with stimulus moment as for the contralateral representation. Bearing in mind the fact that ipsilateral features must be first processed by the contralateral hemisphere, this suggests that around 220 ms, broadly consistent with the tail end of the classical N170 ERP event (see Figure 4), a channel is opened through which features can be transmitted across hemispheres. The N170 has been demonstrated to reflect cross-hemispheric transfer of visual features, with the peak ipsilateral representation of the eyes occurring after the contralateral peak of the N170 event^36^. The linear relationship between stimulus moment and representation moment after this gating event suggests that the channel remains open during the remainder of fixation. Despite the same experimental stimuli, this gating phenomenon is only seen in the gender task, suggesting that it is specific to lateralized task-relevant features (the eyes being used almost exclusively for the gender task). In a recent study, the N170 also appeared to filter out task-irrelevant features: while both task-relevant and task-irrelevant features were processed prior to 170 ms, only task-relevant features were processed afterwards^37^. Of note, the cause of this gating cannot be repetition priming because it is also visible in trials where only one feature is revealed once.

Another notable result is the occurrence of two negative peaks instead of one in the contralateral representation of the eyes in the gender task, with the first one sensitive to only a narrow time window after the stimulus onset. Interestingly, in the case of the right eye, these two peaks have significantly distinct topographies, suggesting distinct neural generators. These generators might resemble the generators of the N170 since the activations are similarly peaking around 170 ms after the reception of eye information. Other studies have observed multiple peaks at the expected timing of the N170^40–41^; these are likely corresponding to activity from different generators. In one study, negative peaks around 160 ms have been found to originate from the fusiform gyrus while negative peaks around 180 ms have been localized as originating from the intraparietal sulcus^41^. Interestingly, if we exclude the first peak and only look at the biggest negative cluster, we notice a pattern that is similar to the positive cluster on the ipsilateral electrodes: all information received in the first ~50 ms is processed at about the same moment (peak around 200 ms) while information received afterwards is processed with a relatively constant (but slightly increasing) offset of 150-170 ms, representation moment increasing with stimulus moment. It is possible that a gating event occurs here too, preventing processing by the sources of this component to start before ~200 ms after the face onset. This gating occurs at about the same latency as the ipsilateral gating, at the expected timing of the classical N170 ERP component.

Other differences in processing across stimulus moments are also visible. For example, the negative activation on ROT has an increased latency for late stimulus moments for some feature/task combinations (that is, this activation peaks after a longer time interval following the reception of information, if this information is received later). This may be a consequence of the prioritization of information received earlier. The visual system is likely to prioritize information received early since it might be unknown for how long information from that stimulus will reach the retina. Thus, the processing of information received late is likely to be delayed or processed more slowly. The opposite phenomenon was visible for the last positive activation in some cases: its latency was greater at early stimulus moments. In other words, there was “temporal compression”: information received earlier was “maintained” for a longer time and all information was processed at almost the same moment independently of when it was received on the retina. It is expected that information received at different moments is processed simultaneously at some point in the brain if it is to be integrated together by higher level areas. The temporal compression we observe may be a consequence of this process of accumulation and integration of information. This is consistent with other studies reporting a component at similar latencies associated with accumulation of evidence and temporal integration^38–39^.

Although adaptation or priming to previously seen features can be ruled out as a source of these differences because they are also present in trials with only one bubble, a bottom-up cause still might have been possible. For instance, different parts of the visual field may always be processed at specific moments during fixation. To investigate whether there were top-down origins to the effects we observed, we verified whether the task modulated them. We found significant interactions between information stimulus moment and task on several sensors at many latencies. In other words, the differences observed in the processing of information received at different moments were not the same depending on the task: consequently, these differences are at least partly top-down in origin. Significant interactions were observed at electrodes and latencies similar to those of the significant effects of stimulus moment but started slightly later, a result that is expected for top-down modulations. Moreover, significant interactions were occurring in slightly different areas. For example, while the processing of the mouth was globally more modulated by stimulus moment on right occipital electrodes, the interaction with the task was stronger on central and left occipital electrodes. This suggests that bottom-up mechanisms and top-down sampling are taking place in different loci.

That the brain processes information differently according to when it was received during fixation, that this occurs even when only one such information is revealed in the course of a trial, and that these differences are modulated by the task, all suggest that each time slot is assigned a different “role” in a top-down fashion. This is compatible with the idea of ballistic visual routines: different operations may be applied to the visual input in a sequential fashion, these operations may vary according to the goal of the computation, and the outcome of the first steps does not change the operations applied thereafter^5,23^. A non-uniform time course of the behavioral use of information in visual recognition has been observed in a few studies^11,14,16^; here, we demonstrate it in the brain for the first time and we show that it is at least partly top-down in origin. Moreover, the variations in processing across stimulus moments relate to variations in behavior; that is, as it could be expected, how the brain (particularly occipito-temporal areas) processes information received at a specific moment relates to how this information will be used to perform the task.

In summary, we uncovered in this study the neural response to specific information received at specific moments during fixation and we showed that when light is received on the retina matters: processing is modulated by the specific moment at which information is received, even within a single eye fixation. These differences can be quite striking, such as an additional delay of 100 ms for information received at some moments. Importantly, these variations remain even when we account for information perceived beforehand, and they are modulated by the task. Moreover, they correlate to differences in the use of information for the task. These results suggest that task-dependent visual routines of information sampling are applied top-down to the continuous visual input.

The novel method introduced in this article also seems a promising avenue to shed light on the accumulation and integration of information occurring during object recognition: indeed, it should allow us to visualize the simultaneous processing, at a given time point and location, of information that was received on the retina at different time points. Future studies using more spatially resolved brain imaging methods such as MEG should investigate how information received at different moments is processed, accumulated, integrated and transferred across brain regions. This method could also be used with intrinsically dynamic stimuli such as dynamic facial expressions or naturalistic movies to investigate how an observer integrates evolving information.

## Acknowledgements

The study was supported by a Discovery Grant (04777-2014) from NSERC to F. G. and by an Alexander-Graham-Bell Doctoral Scholarship from NSERC to L. C.

## Supplementary Figure Legends

**Figure S1.**
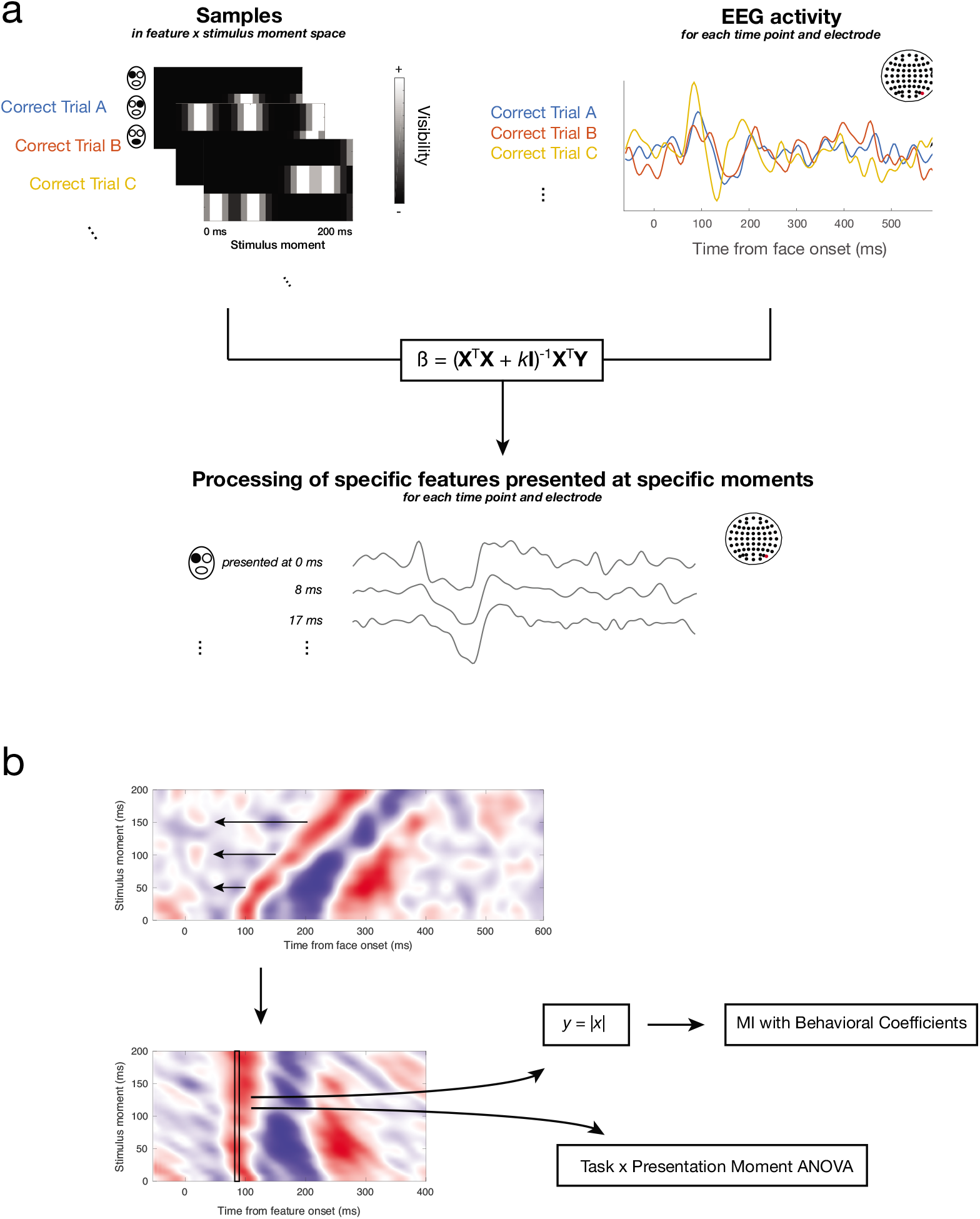
EEG data analyses. **A)** On each trial, a random sampling matrix determines how much each face feature is visible on each presentation moment (the *samples*). Only sampling matrices of truly correct trials (see Methods: EEG Data Analysis) are kept. On each corresponding trial, EEG activity is also recorded across the scalp for a certain period of time (examples are shown for one electrode). For each subject, samples (**X**; independent variable) and EEG activity (**Y**; dependent variable) are combined using a regularized (ridge) multiple linear regression, which allows us to uncover the EEG activity, across time and across the scalp (examples are shown for one electrode), related to the presentation of each specific face feature shown at each stimulus moment. These time courses of regression coefficients can be arranged in images (maps) for specific face features and electrodes where amplitude is now represented by color (see panel B or figures 5 and 6 of the manuscript). **B)** Prior to further analyses, maps of regression coefficients are rearranged so that the zero point is the onset of the feature instead of the whole face (note the change of the *x*-axis title). More specifically, EEG activity related to the presentation of a feature 8.3 ms after the face onset is shifted left by 8.3 ms, EEG activity related to the presentation of a feature 16.7 ms after the face onset is shifted left by 16.7 ms, etc. (see Methods: EEG Data Analysis). Only the first 400 ms are kept so that there is the same number of time points associated with each stimulus moment. Each 24-element column of this realigned image (activity across stimulus moments for each latency from the feature onset) is then submitted to subsequent analyses (example illustrated for one column). In the task x presentation moment ANOVA, columns are compared across subjects and the effect of the task (between-subject factor), the effect of the stimulus moment (within-subject factor), and the interaction between those factors are computed. Prior to the mutual information (MI) analysis, coefficients are transformed into their absolute values. For each subject, mutual information is then computed between the column of values and the vector of 24 values obtained in the behavioral analysis (see Methods: Behavioral data analysis) associated to the same face feature.

**Figure S2.**
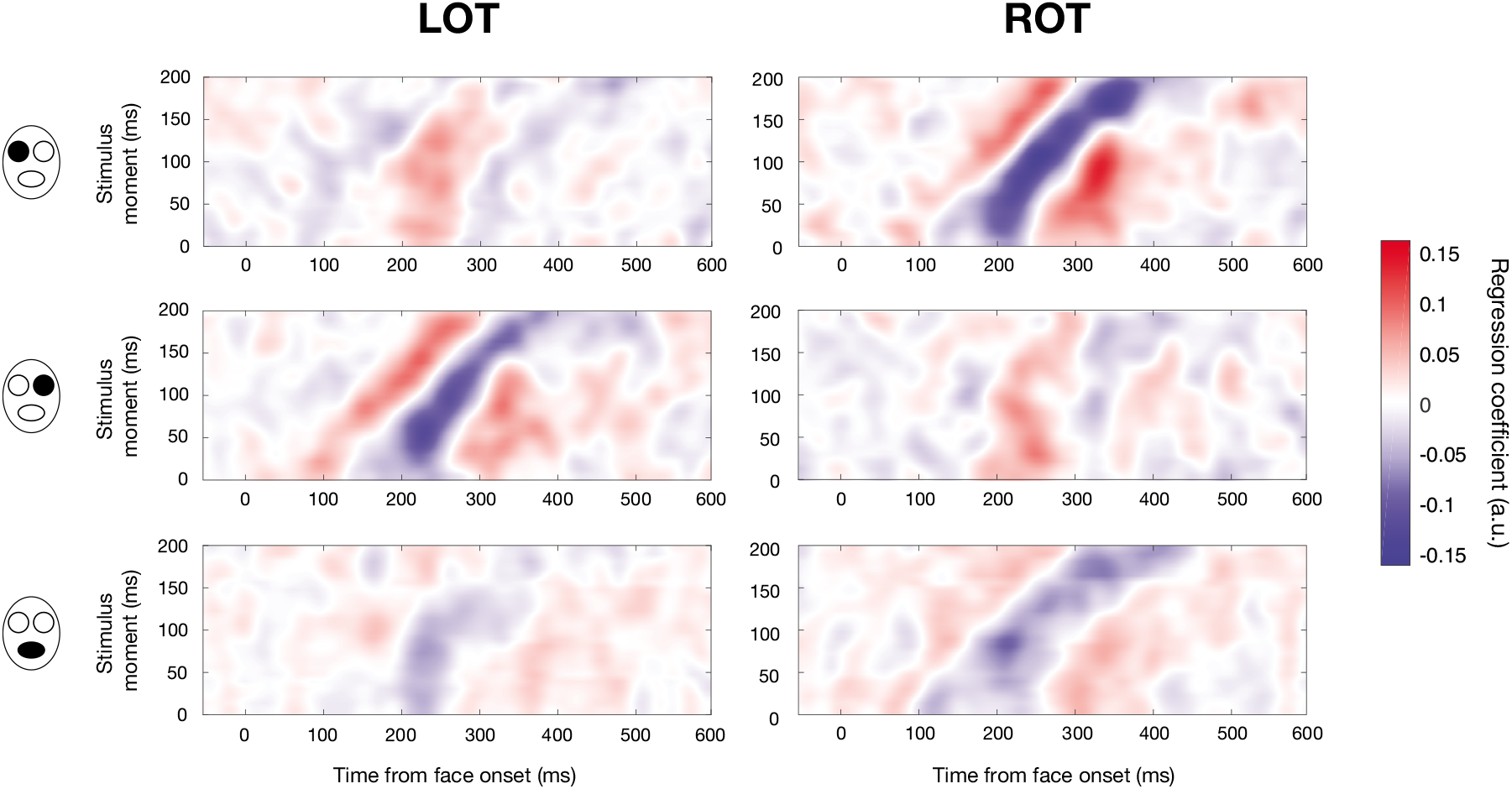
Mean maps of regression coefficients for the gender task, for LOT and ROT sensors (columns) and for each face feature (rows), when including only trials in which there was one bubble (one feature revealed once). See Figure 5 in the main manuscript.

**Figure S3.**
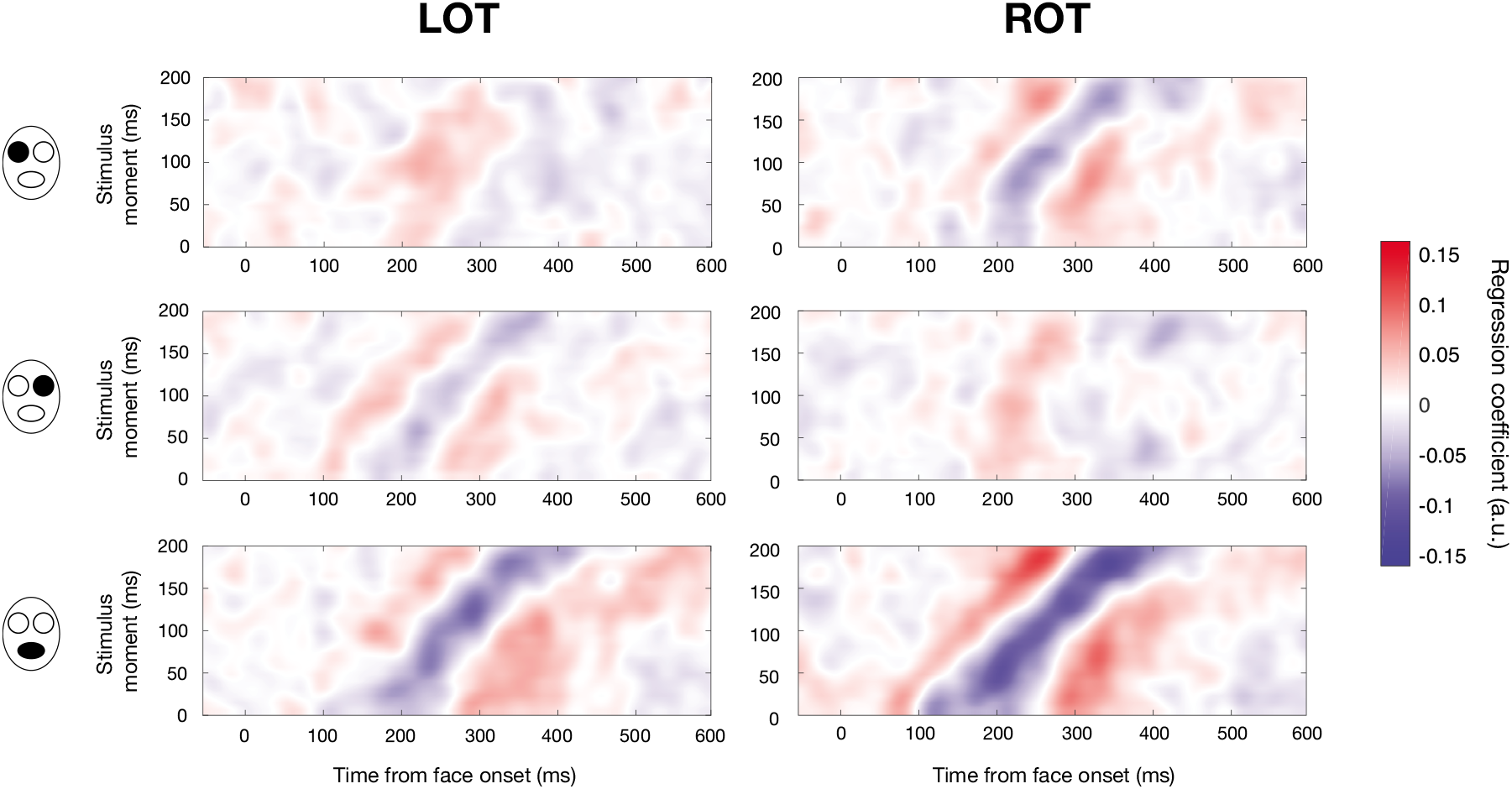
Mean maps of regression coefficients for the expression task, for LOT and ROT sensors (columns) and for each face feature (rows), when including only trials in which there was one bubble (one feature revealed once). See Figure 6 in the main manuscript.

**Figure S4.**
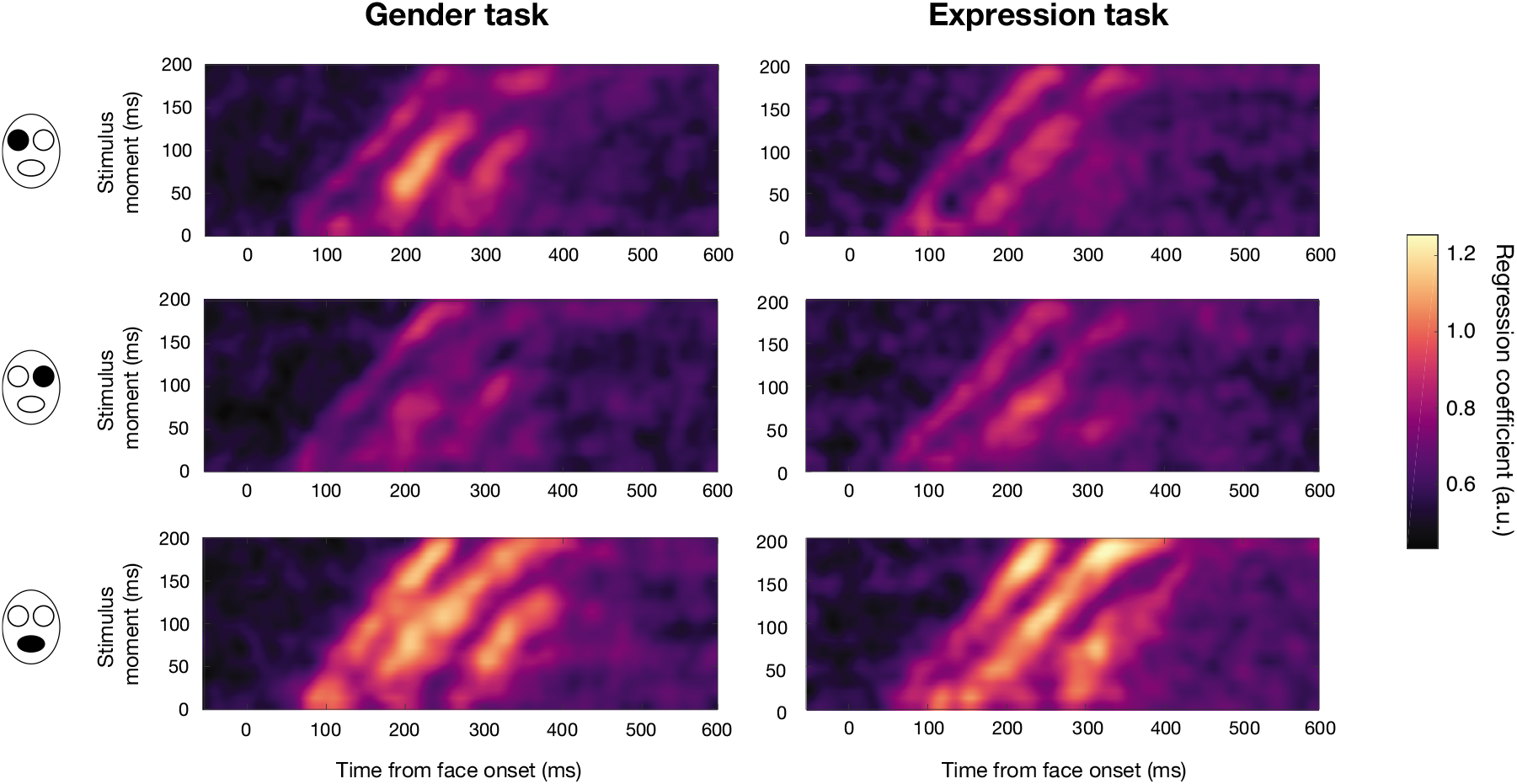
Global scalp regression coefficients for the gender and expression task (columns), for each face feature (rows). To compute these maps, we computed the global field power (standard deviation across sensors) of the regression coefficients for each task and face feature.

## Supplementary Movie Legends

**Movie S1.** Example of a random stimulus.

**Movie S2.** Same as Movie S1; slowed down 10 times.

**Movie S3.** Another example of a random stimulus.

**Movie S4.** Same as Movie S3; slowed down 10 times.

